# A permeable G310 constriction limits K^+^ flux in the activated KCNQ2 pore and is disrupted by the encephalopathy variant G310S

**DOI:** 10.64898/2026.10.05.756556

**Authors:** Shavkatjon Qoshimboyev, Parthiban Marimuthu, Mukhriddin Makhkamov, Jamoliddin Razzokov

## Abstract

KCNQ2 (Kv7.2) subunits contribute to the channels that carry the neuronal M-current, and variants in KCNQ2 cause epilepsies that range from self-limited neonatal seizures to developmental and epileptic encephalopathy (DEE). Structures of KCNQ2 bound to PIP_2_ and cannabidiol show a widened activation gate, yet it has remained unclear which point of the activated pore limits K^+^ flux and how DEE variants at that point act. We simulated the KCNQ2 pore domain (PDB 8J01) in a POPC bilayer under an applied field of about 565 mV, restraining only the terminal residues of S5 and S6 so that G310 and S314 were free. The activated pore conducted K^+^ at 4.3 pS (95% CI 2.5–6.9; 17 net filter crossings in 1.12 µs over four replicas), while the closed-state pore (PDB 7CR3) conducted none in 560 ns. Conduction proceeded by knock-on through a filter that was doubly occupied in 84–92% of frames. Along S6 the narrowest position of the activated pore was not the canonical gate residue S314 but the glycine of the Pro-Ala-Gly motif, G310: activation widens S314 by 3.7 Å but G310 by only 0.7 Å, and restraining G310 to its closed-state width in one replica abolished gate crossings. The DEE variant G310S reduced conduction about six-fold (G310S/wild-type ratio 0.16, 95% CI 0.02–0.66; p = 0.003). Its serine hydroxyls project into the constriction and halve the water column in the G310 plane (8.7 to 4.7 waters), an effect reproduced in CHARMM36m (9.9 to 4.3). The first coordination shell of a K^+^ ion at the constriction was, however, conserved (6.3 vs 6.5 oxygens), because the serine hydroxyls replace the waters they displace. The hydroxyls did not retain the ion: contacts lasted tens of picoseconds, and ions were instead turned back at the serine ring, fewer reaching the gate plane (45 vs 79 excursions per microsecond) and fewer of those crossing it (5% vs 23%). In mixed tetramers carrying one or two variant subunits the effect on the water column was additive, each subunit removing about a quarter of it (f = 0.22 and 0.58 of the full effect). These results identify G310 as the flux-limiting constriction of the activated KCNQ2 pore in our simulations and point to a permeation defect as one component of G310S loss of function.

**Significance:** Loss-of-function variants in the neuronal potassium channel KCNQ2 cause neonatal epilepsy that ranges from self-limited to severe, but for variants that line the pore it is usually not known whether they impair the opening of the channel or the movement of ions through it. Simulating the activated KCNQ2 pore under an applied voltage, we find that K^+^ flux is limited by a glycine of the conserved Pro-Ala-Gly motif that the field’s standard gate metrics do not measure, and that the encephalopathy variant G310S slows permeation roughly six-fold at exactly this point. Its serine hydroxyls take the place of water in the ion’s first shell, yet they do not trap the ion: ions approaching the serine ring are turned back before they cross it. The effect grows in proportion to the number of variant subunits, the situation in the heterozygous channels of patients.

## Introduction

KCNQ2 (Kv7.2) and KCNQ3 (Kv7.3) subunits form the channels that carry the neuronal M-current, a slowly activating, non-inactivating K^+^ conductance that sets the resting potential and damps repetitive firing (Brown and Adams, 1980; Wang et al., 1998; Soldovieri et al., 2011). Variants in KCNQ2 produce a clinical spectrum that runs from self-limited familial neonatal epilepsy to developmental and epileptic encephalopathy (DEE), and the severe end of that spectrum is dominated by missense variants that cluster in the pore module and in the voltage sensor (Soldovieri et al., 2011; Nappi et al., 2024).

Structures have defined the two ends of the gating cycle. The apo channel (PDB 7CR3) has a closed S6 bundle crossing (Li et al., 2021), whereas a structure obtained with phosphatidylinositol 4,5-bisphosphate (PIP_2_) and cannabidiol bound (PDB 8J01) has a dilated gate and was assigned to an open state (Ma et al., 2023). Whether a deposited structure is conductive, however, is a functional question (Trick et al., 2016), and it is usually answered with geometric proxies (Rao et al., 2019): in KCNQ2, cross-subunit distances measured at the residues that line the bundle crossing, principally S314, A317 and L318 (Nappi et al., 2024; Roscioni et al., 2026). These proxies say nothing about a site that lies one helical turn above them.

That site is the glycine of the Pro-Ala-Gly (PAG) motif, G310. In the homologous KCNQ1 channel, Mkrtchyan et al. (2024) showed that the equivalent PAG glycine forms a narrow cavity constriction through which K^+^ passes only after partial dehydration, and proposed that the same constriction may impede transfer in KCNQ2. The prediction has not been tested directly, and G310 does not appear in the distance sets used to classify the KCNQ2 gate.

G310 is also a disease site. The de novo variant G310S causes DEE; it produces a strong loss of function and acts as a dominant negative when co-expressed with wild-type Kv7.2 or with Kv7.3 (Soldovieri et al., 2020; Mosca et al., 2022). Because currents recorded from G310S recover substantially when PIP_2_ synthesis or calmodulin signalling is manipulated, the defect has been interpreted in terms of gating and PIP_2_ dependence (Mosca et al., 2022); whether the permeation pathway itself is affected has not been examined.

Equilibrium simulations have described the KCNQ2 pore gate (Nappi et al., 2024) and a DEE variant in the pore turret (Thompson et al., 2026), but they do not measure permeation. Simulations under an applied field can address both questions, and one such study of KCNQ2, carried out independently of the present work and posted while it was being completed, has now appeared: Roscioni et al. (2026) simulated the closed and the activated pore, together with three gain-of-function variants, and observed K^+^ permeation through the activated pore. Their runs were carried out at 1.1–1.7 V with the bundle-crossing distances restrained, they report no conductance values, and G310 is absent from their analysis. What remains open is therefore (i) how much current the activated KCNQ2 pore carries at physiological-scale voltages, (ii) where along the pathway the flux is limited, and (iii) whether a pore-lining DEE variant impairs permeation itself.

Here we simulate the KCNQ2 pore domain from 8J01 in a lipid bilayer under an applied field of about 565 mV, with G310 and S314 unrestrained, and compare it with the closed-state pore, with a gate narrowed to closed-state width, with a reduced voltage, and with the DEE variant G310S. We find that the activated pore conducts, that the limiting point is G310 rather than the canonical gate, and that G310S suppresses conduction about six-fold: its hydroxyls displace half of the water in the constriction and take its place in the ion’s first shell, and ions are turned back at the serine ring rather than retained by it.

## Materials and Methods

### Systems

The pore module of KCNQ2 (residues G215-K319 of each of the four subunits, corresponding to S5, the pore helix, the selectivity filter and S6) was taken from the cryo-EM structure of the KCNQ2-calmodulin complex with cannabidiol and PIP_2_ bound (PDB 8J01; Ma et al., 2023; Fig. 1A). The same construct boundaries were used by Mkrtchyan et al. (2024) for KCNQ1. Chain termini were capped with acetyl and N-methylamide groups, and 36 heavy atoms missing from the deposited model were rebuilt with PDBFixer (Eastman et al., 2017). The tetramer was embedded in a POPC bilayer (234 lipids, 117 per leaflet) assembled with packmol (Martínez et al., 2009), solvated with 14,982 TIP3P waters (Jorgensen et al., 1983), and neutralised with 138 K^+^ and 137 Cl^−^ (0.51 M with respect to water), giving 83,148 atoms in a box of approximately 9.4 x 9.4 x 9.0 nm. Three additional K^+^ ions were placed at filter sites S1-S3, as in the deposited structure; the initial loading state of the filter is a recognised variable in simulations of K^+^ channels (Domene et al., 2021). The protein was described with Amber14sb (Maier et al., 2015), lipids with Lipid21 (Dickson et al., 2022), and ions with the Joung-Cheatham parameters (Joung and Cheatham, 2008); all charges were the standard, unscaled values.

**Figure 1.**
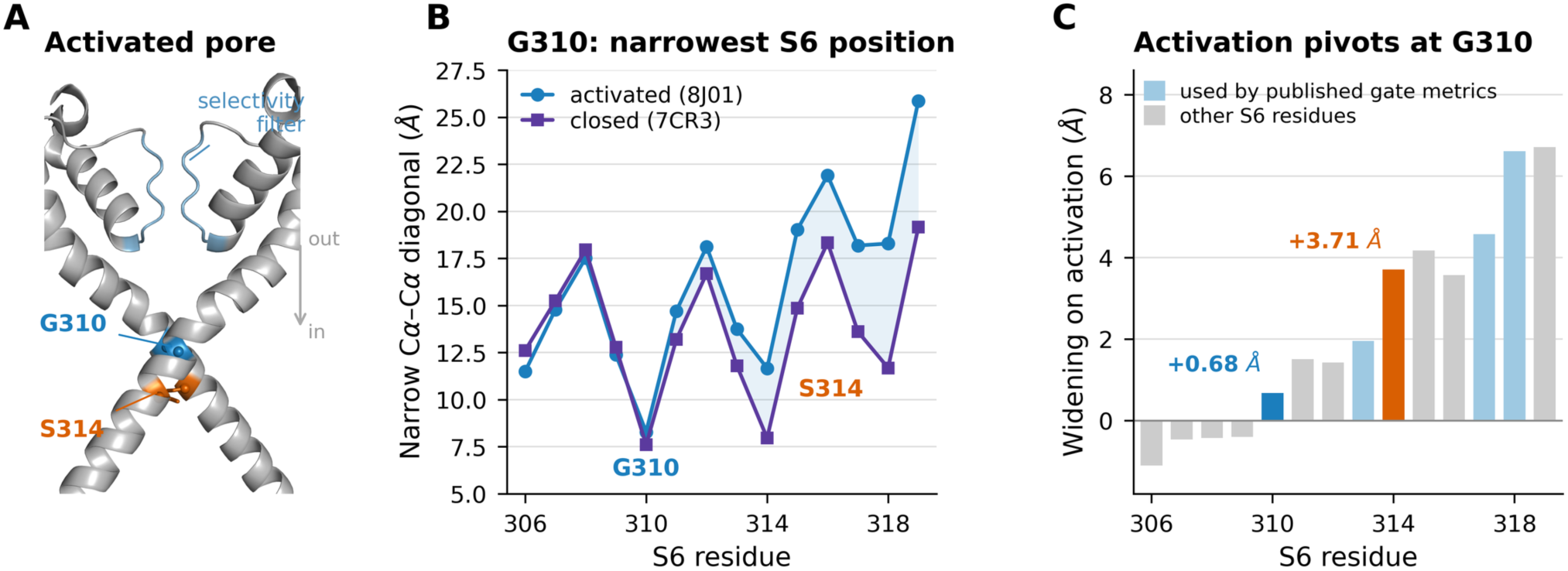
G310 is the narrowest S6 position of the activated pore. **(A)** Two opposite subunits of the activated KCNQ2 pore module (PDB 8J01); the selectivity filter is light blue, G310 blue and S314 orange (spheres, Cα). **(B)** Narrow Cα–Cα diagonal along S6 for the activated (8J01) and the closed (7CR3) structure; G310 is the narrowest S6 position of the activated pore and is close to its closed-state width. **(C)** Widening of each diagonal on activation. S314 opens by 3.71 Å and residues below it by up to 6.7 Å, whereas G310 opens by only 0.68 Å and the residues above it narrow slightly, so activation pivots at G310. Residues used in published gate-distance sets are shaded.

Positional restraints were applied only to the backbone atoms (N, Cα, C, O) of the helix ends that anchor the module in the membrane, residues 215–217 of S5 and 317–319 of S6, with a force constant of 1000 kJ/mol/nm^2^ referenced to the cryo-EM coordinates (keyword POSRES_ENDS). The selectivity filter, G310 and S314 were left free. Of the residues used in published gate-distance sets, A317 and L318 fall within the restrained S6 terminus, so the lower end of the bundle crossing is held near its cryo-EM position; this restraint stands in for the voltage sensor, calmodulin, PIP_2_ and cannabidiol, none of which is present in the simulated system.

Systems were relaxed with a soft-core energy minimisation in which POPC was decoupled (lambda 0.5, 0.3, 0.15, 0.07, 0.03, 0.01) to resolve packing overlaps, followed by standard minimisation, then 0.25 ns NVT (1 fs) and 1 ns NPT (2 fs) with all heavy atoms restrained, 2 ns NPT with backbone restraints, and 5 ns NPT with only POSRES_ENDS.

#### Applied field and production runs

Production runs were 300 ns each with a 2 fs timestep, v-rescale thermostat (Bussi et al., 2007) at 303.15 K (tau 1 ps), C-rescale semi-isotropic barostat (Bernetti and Bussi, 2020) at 1 bar (tau 5 ps), particle-mesh Ewald (Essmann et al., 1995) with a 1.0 nm real-space cut-off, the same cut-off for Lennard-Jones interactions with an analytic dispersion correction, and LINCS constraints (Hess et al., 1997) on bonds to hydrogen. Centre-of-mass motion was removed separately for the protein-lipid and the solvent group. Coordinates were saved every 10 ps.

A constant electric field along z produces a transmembrane potential V = E·L_z_ (Roux, 2008; Gumbart et al., 2012). For each run the field was set to E = 0.565 V / L_z_ of the final equilibration frame, and the potential actually realised, computed as E times the mean L_z_ of the production run, was 549–582 mV across runs (Fig. S1C); conductances are reported with the nominal 565 mV. A second set of runs used 282 mV.

The following systems were simulated (all with identical protocol unless stated; Table S1):

- Wild type (WT; four replicas, 300 ns each). Replicas were started from independent points; velocities were regenerated and each system was re-equilibrated for 2 ns under POSRES_ENDS at zero field before the field was applied.
- G310S (three replicas). The mutation was introduced with PDBFixer into all four chains of equilibrated wild-type frames, the topology was rebuilt, waters overlapping the new side chains were removed (28–36 per system), and the standard equilibration and production protocol was repeated.
- Mixed tetramers (two compositions, two replicas each). Ser310 was introduced into a single subunit (H1) or into two adjacent subunits (H2) of equilibrated wild-type frames, with the same procedure, equilibration and production protocol as for G310S.
- Closed-state pore (two replicas). The pore module of the apo structure (PDB 7CR3; Li et al., 2021) was placed into the equilibrated wild-type environment, with the helix ends restrained to the 7CR3 coordinates.
- Narrowed gate (one replica). Two harmonic distance restraints (GROMACS pull code; k = 15,000 kJ/mol/nm^2^, acting in x and y only) held both G310 Cα–Cα diagonals at 0.761 nm, the value measured in 7CR3, while S314 was left at its 8J01 width.
- Reduced voltage (three replicas at 282 mV).
- CHARMM36m (Huang et al., 2017; two wild-type and two G310S replicas). The Amber systems were converted atom by atom to charmm36-jul2021 with a 1.2 nm cut-off, a force-switch from 1.0 nm and no dispersion correction.
- KCNQ1 benchmark (two replicas of the KCNQ1 pore from PDB 6V01; Sun and MacKinnon, 2020; same protocol).
- Charge-scaled comparison, in which protein charges of Asp, Glu, Lys and Arg and the ion charges were scaled by 0.78 (electronic continuum correction, ECC) following Hui et al. (2025). Two further scaled-charge runs (KCNQ2 and KCNQ1) had both G310 (KCNQ1: G345) Cα–Cα diagonals held at 0.90 and 1.00 nm, respectively, by harmonic distance restraints as in the narrowed-gate control. The starting frames of these two runs were taken from steered simulations in which the diagonals were pulled apart at 0.03 nm/ns with the same harmonic restraint while only the S5 ends were position-restrained; in production the helix-end restraints were referenced to that starting frame rather than to the cryo-EM coordinates, from which the restrained atoms deviated by 1.0 Å (KCNQ2) and 1.2 Å (KCNQ1) r.m.s. All runs of this comparison were analysed like the others, with the first 20 ns discarded: KCNQ2, two scaled-charge and three full-charge runs (one of them 124.5 ns long); KCNQ1, one scaled-charge and two full-charge runs (the second 293 ns long).

Simulations were run with GROMACS 2025.2 (Abraham et al., 2015).

#### Conduction analysis

The primary estimator was the net charge transferred through the selectivity filter. Ion positions were expressed relative to the centre of the filter backbone carbonyl oxygens of residues 276–281, and three zones were defined: the filter (z between −0.8 and +0.45 nm, radius < 0.6 nm), the intracellular cavity (from 0.1 nm above the gate plane to −1.2 nm, radius < 0.9 nm) and the extracellular bulk (z > +1.2 nm). For every filter visit the origin and the destination of the ion were recorded, and the net outward transfer was taken as exits to the extracellular side minus entries from that side. The first 20 ns of every run were discarded, leaving 280 ns per replica. Conductance was computed as G = N*e / (t*V). The three ions placed in the filter at build time were tracked separately from the free ions, since counting them as permeation events is a known source of error.

As an independent check, a displacement estimator was applied to the same trajectories: the signed path length of each ion within the filter segment, divided by the segment length, summed over all ions and frames, together with the equivalent flux through a 0.6 nm slab centred on the gate. This estimator does not use the state machine of the primary method and treats pre-loaded ions as fractional contributions.

For the mixed tetramers the quantity to be read out was fixed before the runs finished: the fraction of the G310S effect on the gate water column, f = (W_WT -W_mixed) / (W_WT -W_G310S), with an additive expectation f = n/ 4 for n variant subunits, and the pre-specified readings were “additive” if the interval contained n/4, “one subunit dominates” if f(H1) was at least 0.5 with 0.25 excluded, and “one subunit is tolerated” if the interval contained 0 but excluded 0.25. Confidence intervals were obtained by the delta method, once from replica standard errors and once from 70 ns block means, and the wider of the two was used. Conduction was designated a secondary measure for these systems, to be reported with its interval rather than read as a conclusion on its own.

Poisson confidence intervals were used for pooled counts, a chi-square test against a Poisson expectation for the consistency of replicas, conditional binomial tests for ratios between systems, and Welch tests for replica-level comparisons.

#### Structural and hydration analysis

Gate geometry was described by the two Cα–Cα diagonals of the four G310 residues, reported as the narrow (d1) and wide (d2) diagonal. The water column at the gate was defined as the number of water oxygens within 0.5 nm of the pore axis and within 0.3 nm of the mean z of the G310 Cα atoms. Cavity occupancy used a 0.6 nm radius between 0.3 nm above the gate plane and −0.8 nm.

The first coordination shell of permeating ions was characterised for every K^+^ within 0.5 nm of the axis by counting water oxygens within 0.30 and 0.35 nm, serine Oγ atoms within 0.35 nm and all other protein oxygens within 0.35 nm. Values at the constriction are reported for ions within 0.25 nm of the G310 Cα plane; bulk reference values were taken from ions more than 3 nm (along z) from the mean z of the lipid phosphorus atoms, i.e. the bilayer midplane, in the same trajectories.

Excursions of ions towards the gate were analysed from the positions of all K^+^ ions within 1.2 nm of the pore axis in every frame of the analysis window. An excursion began when an ion left the intracellular vestibule (more than 0.6 nm below the G310 Cα plane) and entered the lumen (radial distance below 0.6 nm), and ended when the ion either returned below that level or reached the cavity, 0.5 nm above the plane. An excursion was counted as reaching the gate plane if the ion came within 0.25 nm of it, the definition used for the coordination analysis; the deepest position reached and the time spent within the plane were recorded for each excursion, and the outcome is also reported as a function of the depth reached (Table S6B). Contacts with a serine hydroxyl were defined as consecutive frames (10 ps apart) with a K^+^-Oγ distance below 0.35 nm. As a check, visits to the gate plane were counted separately from the ion records of the coordination analysis, which use a different axis definition and independent code. Analyses used MDAnalysis 2.9.0 (Michaud-Agrawal et al., 2011; Gowers et al., 2016).

## Results

### The activated KCNQ2 pore conducts K^+^ under an applied field

Four independent 300 ns replicas of the 8J01 pore module were run at approximately 565 mV with G310 and S314 unrestrained (only the terminal residues of S5 and S6 were restrained; Methods). Counting the net charge carried through the selectivity filter over the analysed 280 ns of each replica gave 5, 5, 4 and 3 outward crossings, or 17 in 1.12 µs in total, corresponding to a conductance of 4.30 pS (95% CI 2.51–6.89; Fig. 2A,B; Table S2). The rate was consistent across replicas (4.30 ± 0.48 pS, mean ± SE; chi-square test against a Poisson expectation, p = 0.89). An independent displacement estimator applied to the same trajectories returned 19.7 net crossings, and a gate-flux estimator 19, agreeing with the primary count to within one event per replica (Fig. S1A).

**Figure 2.**
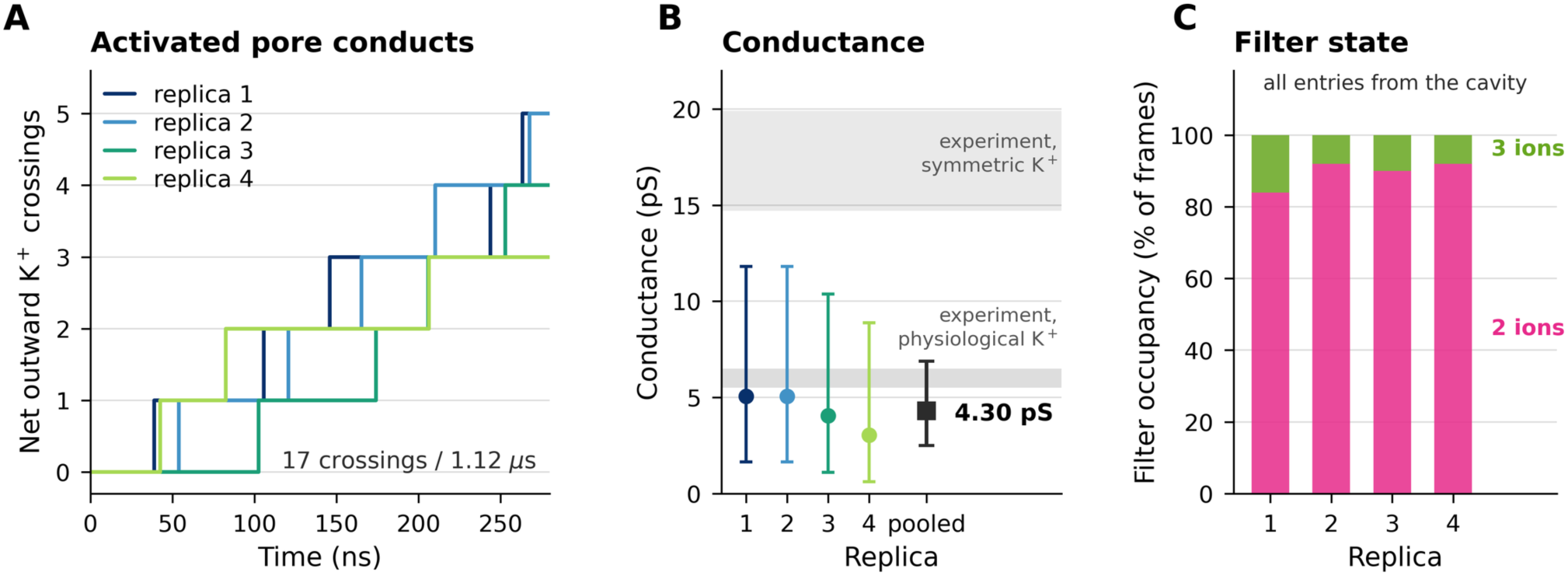
The activated pore conducts K⁺. **(A)** Cumulative net outward K⁺ transfer through the filter in four independent 300 ns replicas at ∼565 mV (first 20 ns discarded): 17 crossings in 1.12 µs; replicas 1–4 are the runs ANE r1, r2, r4 and r5 of Table S2. **(B)** Conductance per replica with Poisson 95 % intervals and the pooled value, 4.30 pS (2.51–6.89); grey bands are single-channel values reported for homomeric KCNQ2 in symmetric and physiological K⁺. **(C)** Filter occupancy: the filter is doubly occupied in 84– 92 % of frames and every entry came from the intracellular cavity, as expected for knock-on conduction.

Twelve of the crossings could be followed as complete single-ion trajectories from the cavity to the extracellular side (Table S2C); the remainder reflect knock-on (Köpfer et al., 2014; Kopec et al., 2018), in which the ion that leaves the filter is not the ion that entered it (Fig. 3A). The filter itself was doubly occupied in 84–92% of frames (Fig. 2C), with transient triple occupancy, and every entry into the filter came from the intracellular cavity, with no entries from the extracellular side. Cl^−^ ions did not enter the pore.

**Figure 3.**
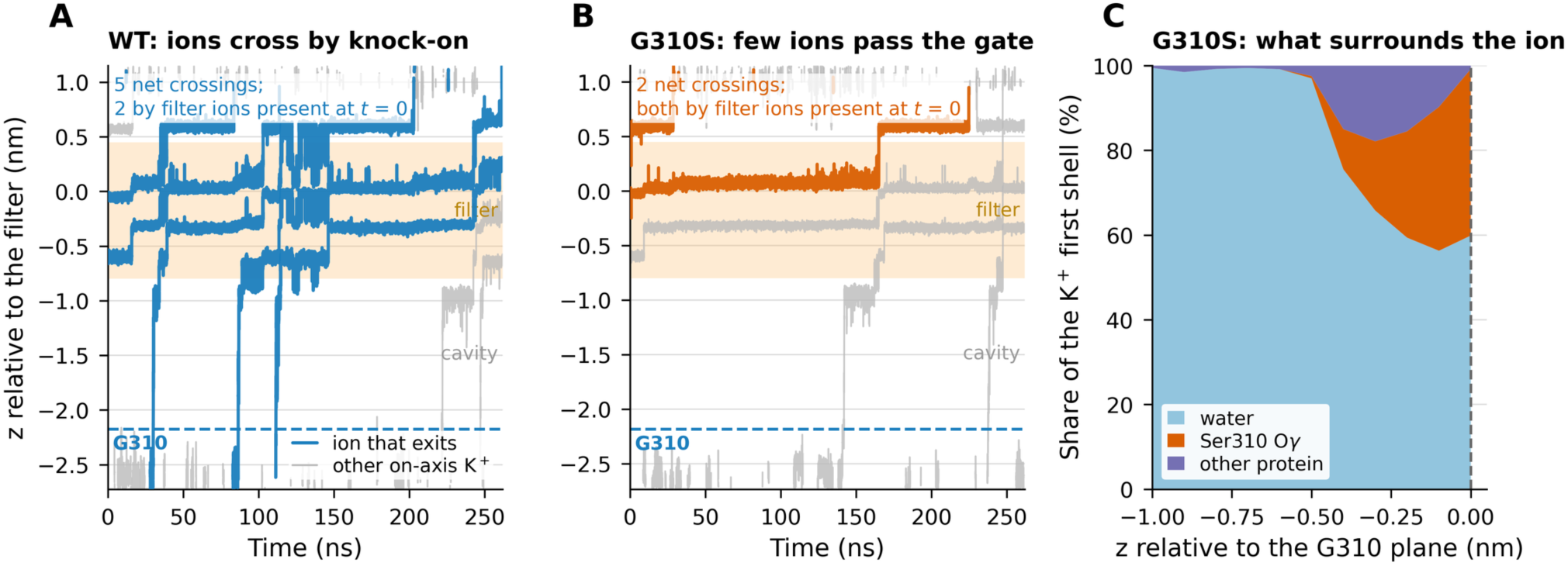
How ions cross the pore in the wild type and in G310S. **(A)** Axial position of every on-axis K⁺ against time in one wild-type replica (ANE r1), relative to the centre of the filter carbonyls; the filter is shaded, the dashed line marks the G310 plane. Ions that eventually leave to the extracellular side are drawn in colour, all other on-axis ions in grey. Ions enter from the cavity, cross G310 and advance through the filter in steps, each entry displacing the ion ahead of it — direct knock-on. Two of the five net crossings are exits of ions that were already in the filter at the start of the window; the other three entered from the cavity. **(B)** The same for the G310S replica with gate crossings (GS r2, the only one of three): ions gather below the gate and only two enter the filter from the cavity; both net crossings are exits of ions that were already inside the filter at the start of the window. **(C)** Composition of the K⁺ first coordination shell in G310S along the axis, as fractions: water is progressively replaced by the serine hydroxyl oxygens as the ion approaches the gate plane, while the total coordination number stays constant (Fig. 5C). Bins with fewer than 30 ion-frames are not drawn, so the profile ends at the gate plane (34 ion-frames at z = 0).

For comparison, single-channel conductances reported for homomeric KCNQ2 are 5.8 ± 0.3 pS (Selyanko et al., 2001), 6.2 ± 0.3 pS (Li et al., 2004) and 6.0 ± 0.2 pS (Hinojo-Perez et al., 2025) under physiological K^+^ gradients, and about 18 pS in symmetric 100 mM K^+^ (Schwake et al., 2000; Fig. 2B). Our simulations use symmetric 0.5 M KCl, so the symmetric measurement is the appropriate reference; the simulated value is about four-fold lower, which is the direction and magnitude expected for fixed-charge force fields (Hui et al., 2025). We therefore treat the absolute conductance as an order-of-magnitude result and base all conclusions on comparisons made within the same protocol.

Applying the same protocol to the KCNQ1 pore (PDB 6V01; Sun and MacKinnon, 2020) gave 11 net crossings in 553 ns, confirming that the protocol transports K^+^ in a second KCNQ channel.

### The closed-state pore does not conduct

Two replicas of the apo pore (7CR3) placed in the same membrane and solvent environment produced no net transfer in 560 ns (ratio to wild type 0, 95% CI 0–0.48; p = 0.001; Fig. 4A,B). One ion entered the filter from the cavity while a filter ion left toward the cavity, so no net charge crossed. The measurement is therefore specific to the activated conformation and is not an artefact of the applied field.

**Figure 4.**
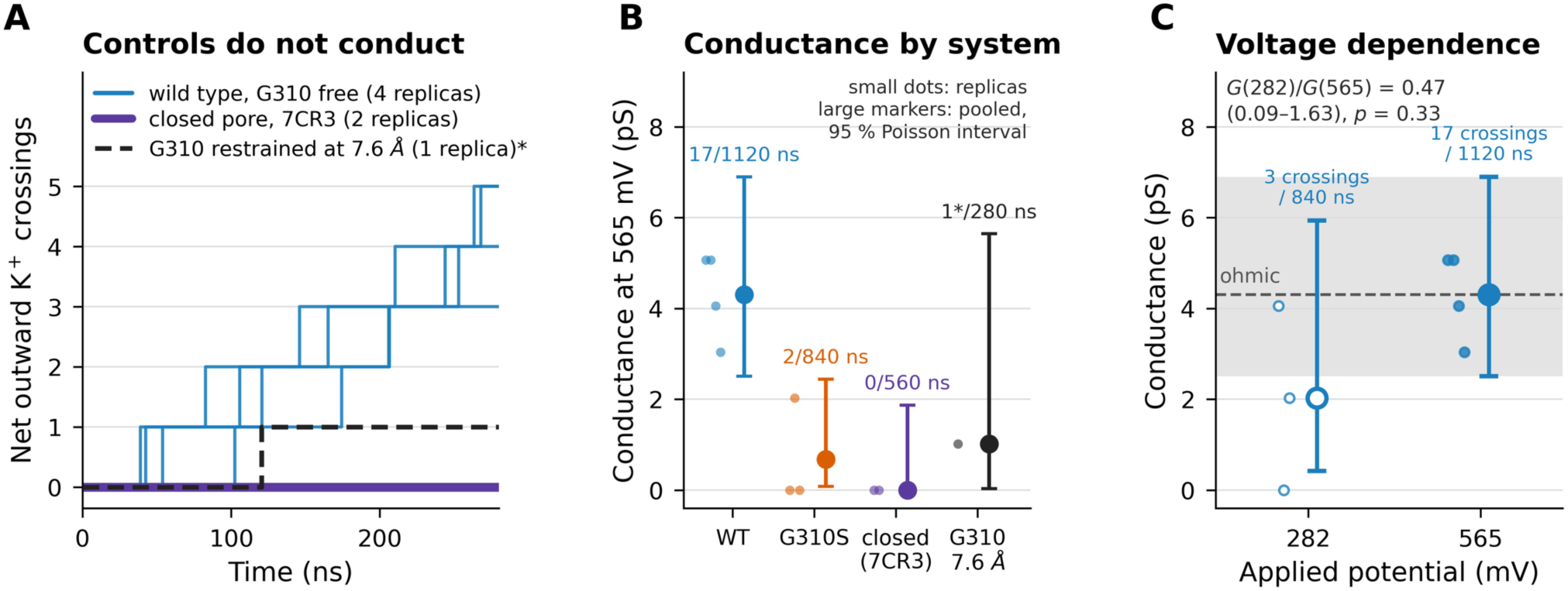
Controls: the closed pore and the narrowed gate do not conduct. **(A)** Cumulative net outward K⁺ transfer for the wild type with G310 unrestrained (four replicas), the closed-state pore (7CR3; two replicas, both at zero) and a pore in which the G310 diagonals were restrained to their closed-state width of 7.6 Å (one replica). Neither control conducts. *The single step in the restrained run is a filter ion that was present at t = 0 leaving; no ion crossed the gate. **(B)** Conductance at ∼565 mV: individual replicas (small dots) and the pooled value of each system (large marker) with its exact 95 % Poisson interval; labels give net crossings / analysed time. **(C)** Conductance of the wild type at 282 mV (open symbol) and 565 mV; small dots are replicas. An ohmic pore has the same conductance at both potentials (dashed line; grey band, 95 % interval of the 565 mV value). G(282)/ G(565) = 0.47 (0.09–1.63, p = 0.33): no deviation from ohmic behaviour is resolved. The 282 mV interval is wide because only three crossings were observed (2, 0 and 1 in the three replicas).

### The flux-limiting point is G310, not the canonical gate

Comparing the activated and the closed structure shows that activation is strongly asymmetric along S6 (Fig. 1B,C). The Cα–Cα diagonal at S314, the residue normally used to describe the KCNQ2 bundle crossing, widens from 7.96 Å in 7CR3 to 11.67 Å in 8J01, an opening of 3.71 Å. At G310, one helical turn above, the same measurement changes from 7.61 Å to 8.28 Å, an opening of only 0.68 Å. In the activated pore G310 is thus the narrowest position along S6, and it remains close to its closed-state width.

A control simulation places the rate-limiting step at this site. When both G310 diagonals were restrained to their closed-state value of 7.61 Å while S314 was left at its 8J01 width, no ion crossed the gate in 280 ns (Fig. 4A), although the same protocol without this restraint gave three to five crossings per replica (4.3 expected; probability of none 0.014). In this single replica a 0.7 Å change at G310 was therefore sufficient to switch the pore off, even though the canonical gate was open.

At the constriction the ion is only mildly dehydrated. A K^+^ at the G310 plane carried 6.09 ± 0.08 water oxygens within 0.35 nm and 0.21 ± 0.03 protein oxygens, against 6.77 water oxygens for K^+^ in bulk solution in the same trajectories (Fig. 5C; Table S4). The ion therefore passes with most of its first shell intact, losing about half a water molecule, which is consistent with the proposal of Mkrtchyan et al. (2024) that the PAG constriction impedes rather than blocks transfer, and rules out an interpretation in which a fully hydrated ion cannot fit.

**Figure 5.**
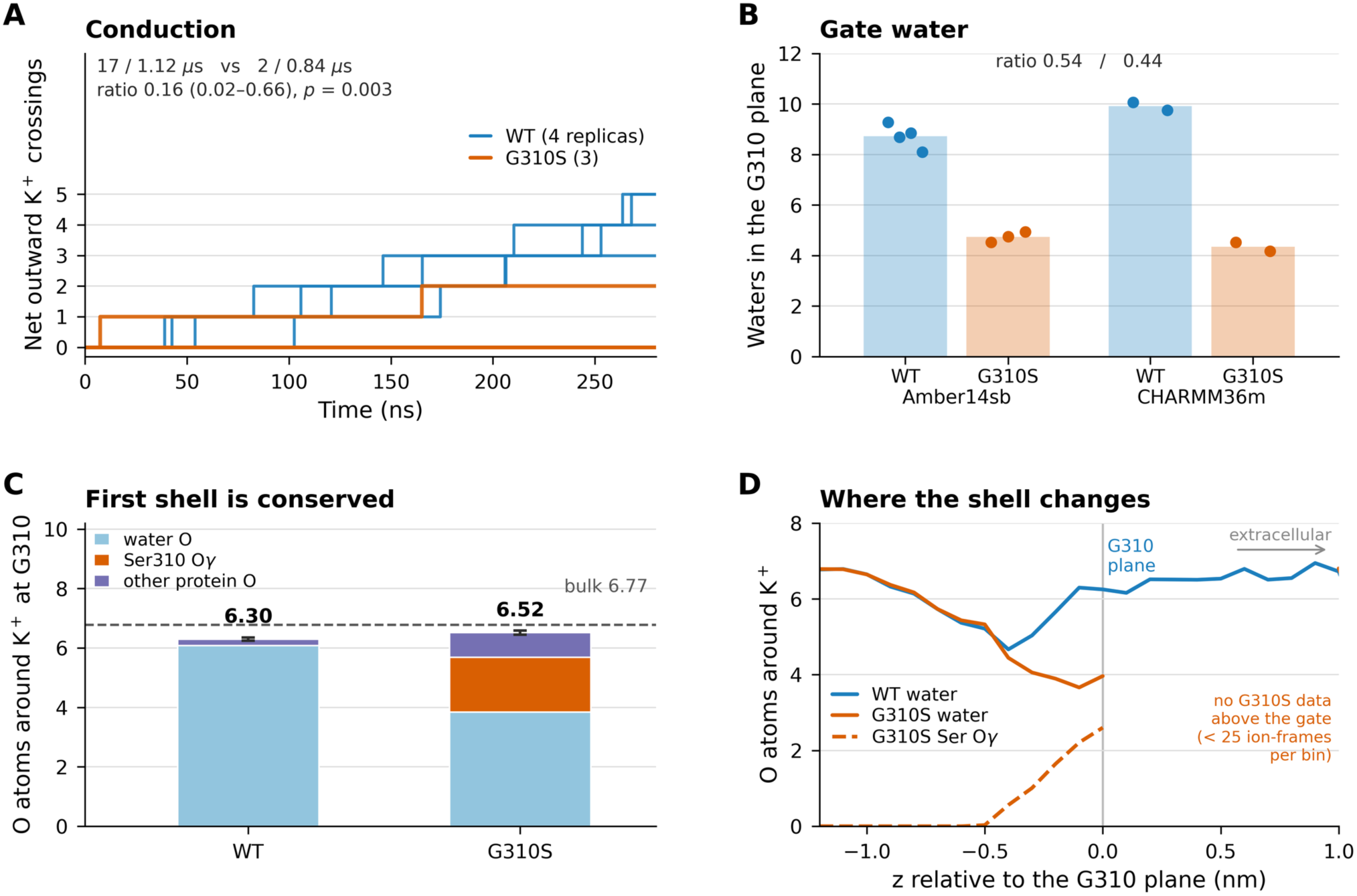
G310S impairs permeation and substitutes hydroxyls for water in the ion’s shell. **(A)** Cumulative net transfer, wild type (4 replicas) versus G310S (3): 17/1.12 µs versus 2/0.84 µs, a ratio of 0.16 (0.02–0.66; p = 0.003). **(B)** Water molecules in the plane of G310; bars are replica means, points individual replicas, in Amber14sb and in CHARMM36m. **(C)** Composition of the first coordination shell of a K⁺ ion at the G310 plane. The variant loses water but gains serine Oγ and other protein oxygens, so the total is unchanged (6.30 ± 0.05 versus 6.52 ± 0.07); the dashed line is the bulk value in the same simulations. **(D)** The same quantities resolved along the pore axis: the two systems are identical until ∼0.4 nm below the gate, where the serine oxygens enter the shell; the G310S curves end at the gate plane because fewer than 25 ion-frames per bin were sampled above it.

### Voltage dependence

Three replicas at 282 mV gave 3 net crossings in 840 ns, or 2.03 pS (Fig. 4C). The ratio of conductances, G(282)/ G(565) = 0.47 (95% CI 0.09–1.63; p = 0.33), is compatible with ohmic behaviour within the statistical resolution of the data; the independent estimators gave 0.61 and 0.70. We detect no deviation from ohmic behaviour and draw no conclusion about rectification.

### The encephalopathy variant G310S suppresses conduction about six-fold

Three replicas of the G310S pore, built by mutating all four subunits in equilibrated wild-type frames and run under an identical protocol, gave 2 net crossings in 840 ns, or 0.68 pS (95% CI 0.08–2.44; Fig. 5A; Table S2). Relative to the wild type this is a ratio of 0.16 (95% CI 0.02–0.66; one-sided conditional binomial p = 0.003); at the level of replicas the difference is also significant (Welch p = 0.013). The independent displacement and gate-flux estimators gave ratios of 0.18 and 0.14 (Fig. S1B), so the effect does not depend on how events are counted. Conduction is reduced roughly six-fold but is not abolished: one replica carried two ions. Both net crossings in that replica were exits of ions already in the filter at the start of the analysis window, displaced by two ions that entered from the cavity, and across all three variant replicas no ion completed a continuous transit from below the filter to above it, against twelve such transits in the wild type (Fig. 3B; Table S2C).

### G310S narrows the water column without narrowing the pore

The variant changes the pore geometry only marginally. The narrow G310 Cα–Cα diagonal contracts from 8.39 Å to 8.17 Å (p = 0.004), a difference of 0.2 Å that is an order of magnitude smaller than the 3.7 Å of gate opening. What changes is the solvent. Water occupancy of a narrow constriction is the usual measure of hydrophobic gating (Beckstein and Sansom, 2003; Jensen et al., 2010; Trick et al., 2016; Gu and de Groot, 2023), and it can be predicted from pore radius and hydrophobicity (Rao et al., 2019); we counted it in the plane of the gate. The serine hydroxyl oxygens sit 3.53 Å from the pore axis, inside the lumen, and the number of water molecules in the plane of the gate falls from 8.73 ± 0.24 to 4.73 ± 0.12 (Welch p = 9e-5; Fig. 5B). Neither the water content of the cavity above the gate (23.4 vs 23.1 waters) nor the delivery of ions to the gate from below (K^+^ present in 15.8% vs 15.0% of frames) changed, so the variant acts inside the constriction rather than by starving it of ions or water. Occupancy of the cavity itself was lower in the variant (5.8% vs 1.3% of frames) but with a wide spread across replicas (p = 0.10), and we do not draw a conclusion from it.

### The ion’s first shell is conserved: serine hydroxyls replace the displaced water

Because the variant halves the water in the gate plane, one might expect the permeating ion to be more strongly dehydrated there. It is not. At the G310 plane a K^+^ ion in the variant was surrounded by 3.85 ± 0.08 water oxygens, down from 6.09 ± 0.08 in the wild type (p < 1e-4), but it simultaneously gained 1.84 ± 0.11 serine Oγ oxygens (p = 0.003) and 0.83 ± 0.13 other protein oxygens (p = 0.03). The total number of oxygens in the first shell was therefore 6.52 ± 0.07 in the variant against 6.30 ± 0.05 in the wild type, a difference that is marginal and in the opposite direction (p = 0.06; Fig. 5C; Table S4). Resolved along the axis (Figs. 3C and 5D), the two systems are indistinguishable until about 0.4 nm below the gate, at which point the serine oxygens begin to enter the ion’s shell and the water count diverges, with the substitution complete at the gate plane itself. The bulk reference value, 6.77 waters, was identical in both sets of simulations.

The variant therefore does not strip the ion of its solvation shell; it replaces a labile part of that shell with oxygens attached to the protein.

### Ions are turned back at the serine ring, not retained by it

If part of the ion’s shell is supplied by oxygens tethered to the protein, the ion might be expected to dwell at the serine ring. We therefore followed every ion that left the intracellular vestibule towards the gate, from 0.6 nm below the G310 plane until it either returned below that level or arrived in the cavity, 0.5 nm above the plane (Methods). In the wild type 88 such excursions reached the gate plane in 1.12 µs (79 per us) and 20 of them continued into the cavity (23%); in the variant 38 reached it in 0.84 µs (45 per us; rate ratio 0.58, p = 0.005) and 2 continued (5%; Fisher p = 0.02; Table S6A). The loss of crossings (20 against 2; rate ratio 0.13, compared with 0.16 for the net filter count) is therefore the product of fewer arrivals at the plane and a lower probability of passing it.

The ion was not held at the ring. The time spent in the gate plane per excursion was no longer in the variant than in the wild type (mean 117 against 141 ps, medians 20 against 60 ps; Welch p = 0.26 on replica means), the plane was occupied for 0.53% of the time against 1.11%, and individual contacts between an ion and a serine Oγ lasted a median of 30 ps, with 90% shorter than 0.2 ns and the longest 1.6 ns. Resolving the excursions by the depth they reached locates the loss (Table S6B). Down to 0.25 nm below the plane the variant differed from the wild type by less than two-fold (ratio of rates 0.57–0.73), but only 2 of the 38 excursions that came that far went on to the centre of the plane, against 32 of 89 in the wild type; both ions that reached the centre in the variant crossed. Ions are thus turned back in the lower half of the serine ring, the 0.25 nm over which the hydroxyl oxygens enter the first shell and in which the water column is halved, rather than being trapped once they are coordinated. Counting visits to the gate plane from the ion records of the coordination analysis, with a different axis definition and independent code, gave the same crossing counts (20 and 2) and likewise no lengthening of the dwell time.

### Mixed tetramers: the water column scales with the number of variant subunits

Patients carrying G310S are heterozygous, so their channels assemble with one to three variant subunits. We therefore simulated two mixed pores under the identical protocol, one carrying Ser310 in a single subunit (H1) and one carrying it in two adjacent subunits (H2), with two 300 ns replicas each. The quantity to be read out, and the additive expectation against which to read it, were fixed before these runs finished (Methods).

The water column in the gate plane fell from 8.73 ± 0.24 waters in the wild type to 7.83 ± 0.06 in H1 and to 6.43 ± 0.41 in H2, against 4.73 ± 0.12 in the full variant. Expressed as the fraction f of the G310S effect, this is f = 0.22 (95% CI 0.13–0.32) for one serine and f = 0.58 (0.37–0.78) for two, against additive expectations of 0.25 and 0.50 (Fig. S3A,B; Table S5). Both intervals contain the additive value, so each variant subunit removes about a quarter of the water column independently of the others. Recomputing the same quantity with a different definition of the pore axis and a different set of frames gave 0.24 and 0.60. The narrow G310 diagonal followed the same pattern (8.40 Å in H1 and 8.22 Å in H2, against 8.39 Å in the wild type and 8.17 Å in G310S), whereas the serine hydroxyl pointed less far into the lumen when it stood alone (hydroxyl oxygen 4.69 Å from the pore axis in H1, 4.02 Å in H2 and 3.53 Å in G310S).

Conduction, which was designated the secondary measure for these systems, was not detectably reduced in either mixed pore: 7 net crossings in 560 ns in each, a ratio to the wild type of 0.82 (95% CI 0.29–2.09) in both cases, against 8.5 crossings expected at the wild-type rate and 1.3 at the G310S rate. Compared with the full variant the mixed pores carried about five times the rate (5.3, 95% CI 1.0–52; one-sided p = 0.025). Continuous cavity-to-exterior transits, counted with the independent tracker used in Table S2C, occurred three times in H1 and four times in H2 in 560 ns each, against twelve in 1.12 µs in the wild type and none in 840 ns in G310S (Table S5). These counts are too few to place the mixed pores on the conductance scale between the two homotetramers, and the structural measure is the one that answers the question.

### The structural effect is reproduced in a second force field

Because absolute conduction in potassium channels is known to depend on the force field (Furini and Domene, 2020; Ocello et al., 2020), the wild-type and variant systems were converted atom by atom to CHARMM36m and run for 4 x 300 ns. Under CHARMM36m the wild-type pore did not conduct: no ion entered the filter from the cavity in 560 ns, against the 8.5 entries expected at the Amber14sb rate (p = 0.001; Fig. S2A; Table S3), and the block lay at the filter, not at the gate, since the cavity was occupied in 13–21% of frames (Fig. S2B). This reproduces a known sensitivity of filter occupancy and knock-on mechanism to the force field (Mendoza Uriarte and Roux, 2026).

The structural signature of the variant, however, transferred. In CHARMM36m the gate water column fell from 9.91 to 4.34 waters (ratio 0.44, against 0.54 in Amber14sb; Fig. 5B, Fig. S2C, Table S3) and the serine Oγ atoms sat 3.33 Å from the axis, essentially as in Amber14sb. The mechanism by which G310S acts is therefore not an artefact of one force field, even though the absolute conductance is model-dependent.

A related control used charge scaling of the ionic and charged-residue charges by 0.78. In structure- and width-matched pairs of the KCNQ2 and KCNQ1 pores this reduced net transfer about eight-fold (2 crossings in 840 ns against 23 in 1.22 µs; ratio 0.13, 95% CI 0.01–0.51, p = 3.2e-4) and held the filter at three ions in 97% of frames (Fig. 6A,B). A KCNQ2 run with the gate restrained 0.55 Å wider than in the full-charge runs conducted at a similar rate (7 crossings in 280 ns against 12 in 664.5 ns; Fig. 6C), which suggests that scaling shifts the gate width at which conduction appears rather than abolishing it; this rests on single runs. The result extends to KCNQ channels a caveat already raised for this combination of scaled charges with Amber14sb (Hui et al., 2025); the dependence of permeation on charge strength has also been examined for MthK (Lam and de Groot, 2023).

**Figure 6.**
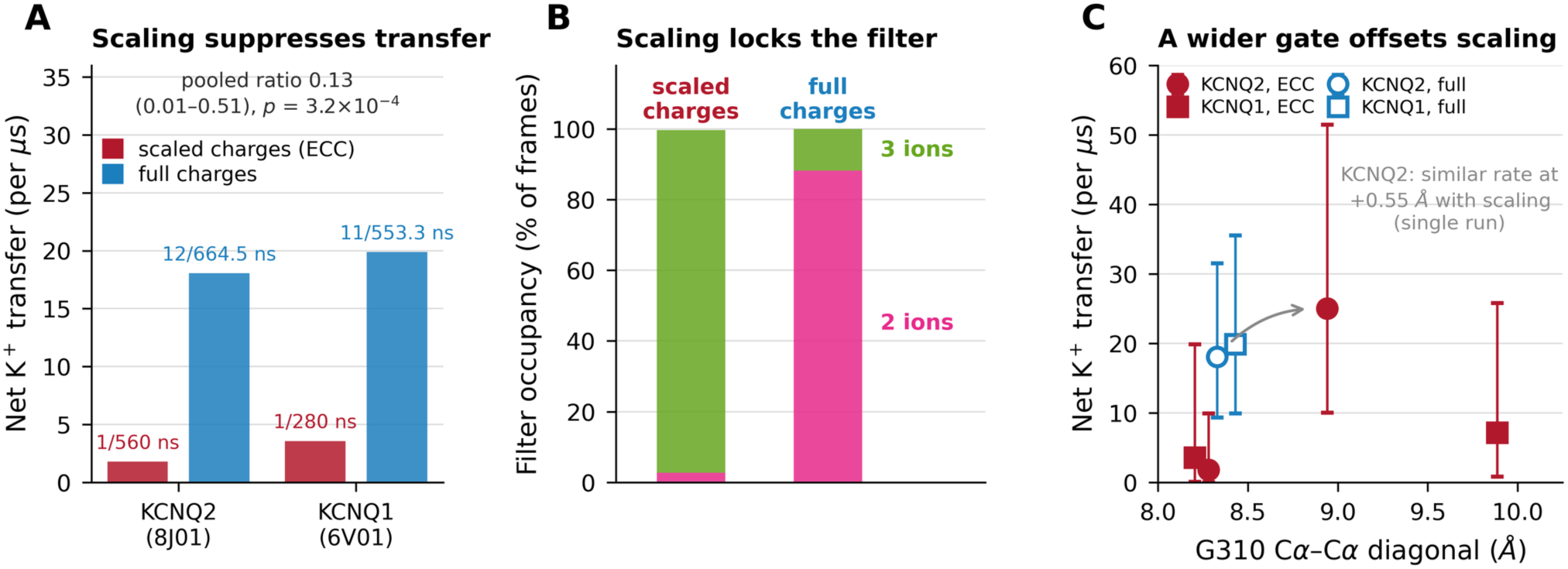
Charge scaling suppresses transfer, and a wider gate partly offsets it. **(A)** Net K⁺ transfer with scaled (Amber14sb-ECC, q = 0.78) and with full charges, for structure -and width-matched pairs of the KCNQ2 and the KCNQ1 pore: 2 crossings in 840 ns against 23 in 1.22 µs, a ratio of 0.13 (0.01–0.51; p = 3.2×10⁻⁴); as elsewhere, the first 20 ns of every run were excluded. **(B)** Filter occupancy: with scaled charges the filter holds three ions in 97 % of frames, whereas with full charges it is doubly occupied in 88 %. **(C)** Net transfer against the narrower G310 diagonal, with exact 95 % Poisson intervals; each point is one run or the pool of one system’s runs, and the two full-charge points are offset horizontally by ±0.06 Å for legibility. In KCNQ2 a scaled-charge run with the gate restrained 0.55 Å wider than in the full-charge runs reached a rate similar to theirs (7 crossings in 280 ns against 12 in 664.5 ns); in KCNQ1 the scaled-charge run with a gate restrained 1.5 Å wider (2 crossings in 280 ns) has an interval that overlaps the full-charge one. The width shift is thus an estimate from single runs, not a measured threshold.

## Discussion

Two results follow from these simulations. The activated KCNQ2 pore conducts K^+^ under an applied field, and the point at which the flux is limited is the glycine of the PAG motif, G310, rather than the bundle-crossing residues normally used to decide whether a structure is open. The encephalopathy variant G310S acts precisely at this point and reduces conduction about six-fold.

### G310 and the description of the KCNQ2 gate

Activation opens S314 by 3.7 Å and G310 by 0.7 Å. A structure can therefore satisfy every criterion applied to the bundle crossing while retaining a constriction one turn above it, and the geometric sets used to classify the KCNQ2 gate, which measure S314, A317 and L318 (Nappi et al., 2024; Roscioni et al., 2026), do not register that site. Restraining G310 to its closed-state width, which changes the pathway by 0.7 Å and leaves S314 untouched, was sufficient to stop conduction in the replica we ran. The sensitivity of the conductance to this residue argues that G310 belongs in the description of the KCNQ2 gate.

The result extends to KCNQ2 the mechanism that Mkrtchyan et al. (2024) established for KCNQ1, where the equivalent PAG glycine forms a cavity constriction through which ions pass after partial dehydration, and where they suggested that the same feature might impede transfer in KCNQ2. Our measurements support the weaker of the two possible readings of that suggestion: the constriction impedes but does not block. The ion loses about half a water molecule from its first shell as it crosses, so an argument based on the radius of a fully hydrated K^+^ does not describe what happens, as has also been found for a constricted opening in Kir channels (Black et al., 2020).

### Relation to concurrent work on the same structure

Independently of the present work, and posted while it was being completed, Roscioni et al. (2026) reported permeation through the activated 8J01 pore, together with three gain-of-function variants, using applied potentials of 1.1–1.7 V with the S314 and A317 distances restrained. Their observation that the activated pore conducts is consistent with ours. The two studies differ in what they were designed to measure: they examined how gain-of-function variants stabilise a partially open gate and state explicitly that their runs were not intended to yield conductance values, whereas we measured conductance at physiological-scale voltages, tested a loss-of-function variant, and located the limiting site. Both studies hold the lower end of S6 in place, theirs through the S314 and A317 distances and ours through positional restraints on residues 317–319, which leave S314 and G310 free. G310 does not appear in their analysis. The two sets of results are complementary, and one of their observations supports ours: they too found that the conduction behaviour of this pore differs between CHARMM36m and Amber force fields.

### What kind of defect G310S is

The variant places one hydroxyl group per subunit into the narrowest part of the gate region and halves the water column there, yet the ion that crosses is coordinated by the same number of oxygens as in the wild type, because the serine hydroxyls take the place of the waters they exclude. One reading of such a result would be that the serine ring acts as a binding site that holds the ion, its tethered oxygens exchanging more slowly than water. The trajectories do not support that reading: contacts between ion and hydroxyl last tens of picoseconds, the ion spends no longer in the gate plane than in the wild type, and the plane is occupied half as much of the time. What changes is the outcome of the approach. Ions arrive at the plane somewhat less often and, having arrived, cross it about four times less often, and the loss occurs in the lower half of the ring, where the hydroxyl oxygens enter the shell and the water column that would carry the ion onwards is depleted. G310S is thus better described as a defect of entry into the constriction than as a trap within it. In a channel that conducts by knock-on (Köpfer et al., 2014; Kopec et al., 2018; Lam and de Groot, 2023), in which every ion entering the filter comes from the cavity, a lower probability of passing the constriction translates directly into a lower current.

We have not determined why the ion is turned back. A six-fold reduction in rate corresponds to only about 4.7 kJ/ mol if it arises from an equilibrium barrier, and an energetic origin of that size is neither demonstrated nor excluded by our data. Because conduction in these channels is thought to depend on the relative free energies of multi-ion states (Hui et al., 2025), separating a small increase in the barrier from an effect of the interrupted water column on the motion of the ion would require multi-ion free-energy calculations or longer sampling of the constriction, and is the natural next test.

### Consequences for the G310S phenotype

G310S is a strong loss-of-function variant that acts as a dominant negative and forms heteromers with Kv7.3 (Soldovieri et al., 2020; Mosca et al., 2022). Currents from G310S recover when PIP_2_ synthesis or calmodulin signalling is manipulated, which has been read as a gating defect (Mosca et al., 2022). Our results do not contradict that reading; they add a second, independent defect. Whole-cell current is the product of single-channel conductance, open probability and the number of channels at the surface. Taking our conductance ratio of 0.16, and the 5.2-fold increase in surface expression reported for G310S, the measured current ratios are arithmetically consistent with a channel that both conducts poorly and opens poorly, with the opening defect largely relieved under maximal PIP_2_ and calmodulin stimulation while the conductance defect persists. This is a consistency argument, not a measurement, and it makes a direct prediction: single-channel recordings of G310S under conditions that restore its open probability should show a unitary conductance near 1 pS rather than the 6 pS of the wild type.

Our simulations describe a homotetramer carrying four serines, whereas patients are heterozygous and their channels carry one to three variant subunits, often together with Kv7.3. To ask how the defect scales with that number, we simulated pores with one and with two adjacent variant subunits. The structural effect is additive: each serine removes about a quarter of the water column in the gate plane, with no indication that a single subunit dominates the effect or that it is tolerated (Fig. S3; Table S5). The homotetramer is therefore the limiting case of a graded defect rather than a state of its own, and a channel carrying one or two variant subunits should present an intermediate constriction. Conduction in the mixed pores was not measurably reduced, but with seven events per 560 ns their rate is bounded only loosely, and we do not infer from it how the rate itself scales with subunit number. Additivity of the structural defect does not by itself account for the dominant-negative behaviour reported in expression systems, which concerns the current carried by a whole population of channels and may also reflect assembly and gating.

### Limitations

The number of permeation events is small: 17 in the wild type and 2 in the variant, so the confidence interval on the suppression factor spans roughly 1.5-fold to 60-fold, and the conclusion rests on the ratio rather than on the individual conductances. Absolute conductance is force-field dependent (Furini and Domene, 2020; Ocello et al., 2020; Mendoza Uriarte and Roux, 2026), and under CHARMM36m the wild-type pore did not conduct at all, with the block located at the filter; the structural effect of the variant, however, transferred between the two force fields. The mixed tetramers were run as two replicas per composition, and their event counts constrain the conduction rate only loosely; the additivity conclusion rests on the water column, which is the measure that was fixed in advance for that purpose. The applied potential is several-fold above physiological, the ionic strength is 0.5 M, and the variant was simulated only in the activated conformation, so nothing here addresses whether G310S also alters gating. The narrowed-gate control is a single replica, and the charge-scaling comparison rests on single runs at each gate width. The excursion analysis rests on two crossings in the variant, so the probability of crossing is bounded only loosely, and the fraction of arrivals that cross depends on the depth at which an arrival is counted (Table S6B). The pore module was simulated without the voltage sensor, calmodulin, PIP_2_ or cannabidiol, and its activated conformation was maintained by restraints on the S5 and S6 termini, which include A317 and L318; the simulations therefore do not address the stability of the open state. Finally, the study is computational; the prediction above is the experiment that would test it.

## Author contributions

S.Q.: conceptualization, methodology, software, formal analysis, investigation, data curation, visualization, writing -original draft. P.M.: supervision, writing -review and editing. M.M.: supervision, writing -review and editing. J.R.: supervision, resources, writing -review and editing.

## Competing interests

The authors declare no competing interests.

## Funding

This research received no specific grant from any funding agency in the public, commercial or not-for-profit sectors.

## Data and code availability

Simulation input files for every system in Table S1 (GROMACS topologies, parameter, coordinate and index files), the analysis and figure scripts, the per-run event tables behind Table S2 and the processed data behind every figure are available at https://github.com/Kushimboev/kcnq2-g310. Full production trajectories (about 9.3 GB per 300 ns run, approximately 190 GB in total) are too large for that repository and are available from the corresponding author on request. Reduced trajectories containing the protein backbone, the pore-lining residues and all ions, which are sufficient to reproduce every conduction count reported here, will be deposited on Zenodo together with an archived release of the repository.

## Use of artificial intelligence tools

A large language model (Claude, Anthropic) was used under the authors’ direction to write and refine the analysis and figure-generation code and to draft and edit the text of this manuscript. It was not used to generate, select or alter any simulation or measurement result. The authors verified the code and the numbers independently, reviewed all text, and take full responsibility for the content.

## Supplementary Information

### Supplementary Tables

**Table S1.** Simulated systems.

| System | What was simulated | Replicas | Analysed time | Potential |
| --- | --- | --- | --- | --- |
| Wild type | Pore module of the deposited activated structure (PDB 8J01), full ionic charges | 4 | 4 × 280 ns | ~565 mV |
| G310S | Serine at position 310 in all four chains, introduced into equilibrated wild-type frames | 3 | 3 × 280 ns | ~565 mV |
| Closed-state pore | Pore module of the apo structure (PDB 7CR3) placed in the equilibrated wild-type environment | 2 | 2 × 280 ns | ~565 mV |
| Narrowed gate | Wild type with both G310 Ca–Ca diagonals restrained at 0.761 nm | 1 | 280 ns | ~565 mV |
| Reduced potential | Wild type | 3 | 3 × 280 ns | 282 mV |
| CHARMM36m | Wild-type and G310S systems converted atom by atom to CHARMM36m | 2 + 2 | 4 × 280 ns | ~565 mV |
| KCNQ1 benchmark | Pore module of KCNQ1 (PDB 6V01), same protocol | 2 | 280 + 273 ns | ~565 mV |
| Charge scaling (ECC, q = 0.78) | Scaled-charge runs of the KCNQ2 (2) and KCNQ1 (1) pores with matched full-charge runs (KCNQ2 3, KCNQ1 2); two further scaled-charge runs with the G310 diagonals restrained wider (KCNQ2, KCNQ1) | 3 + 5 + 2 | 840 ns scaled, 1218 ns full-charge, 2 × 280 ns wider gate | ~565 mV |
| Mixed tetramers | Ser310 in one subunit (H1) or in two adjacent subunits (H2), introduced into equilibrated wild-type frames | 2 + 2 | 4 × 280 ns | ~565 mV |

**Table S2.**
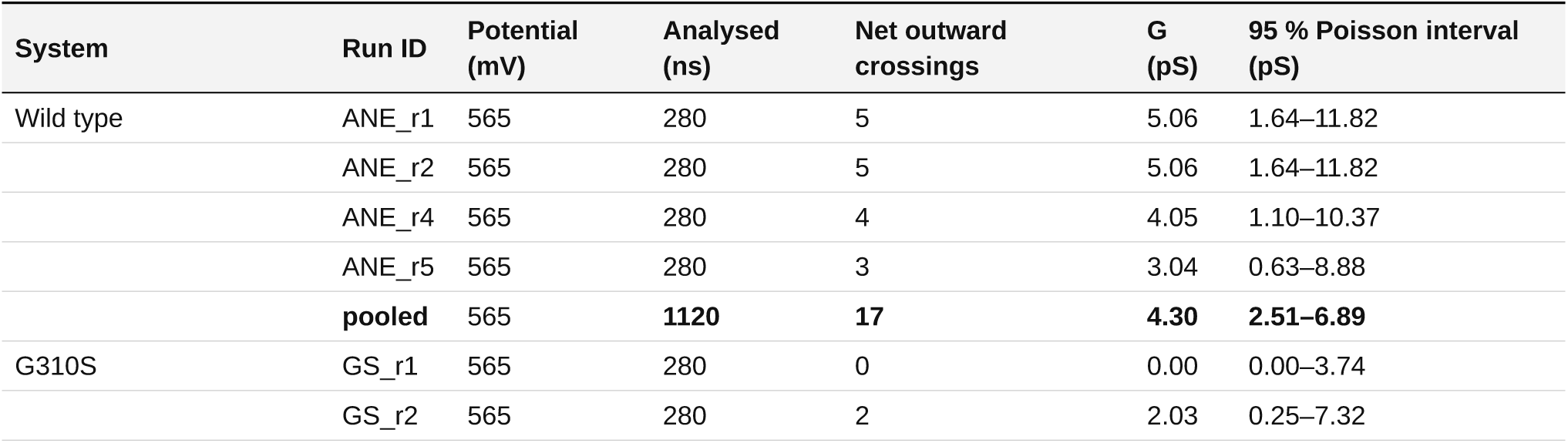
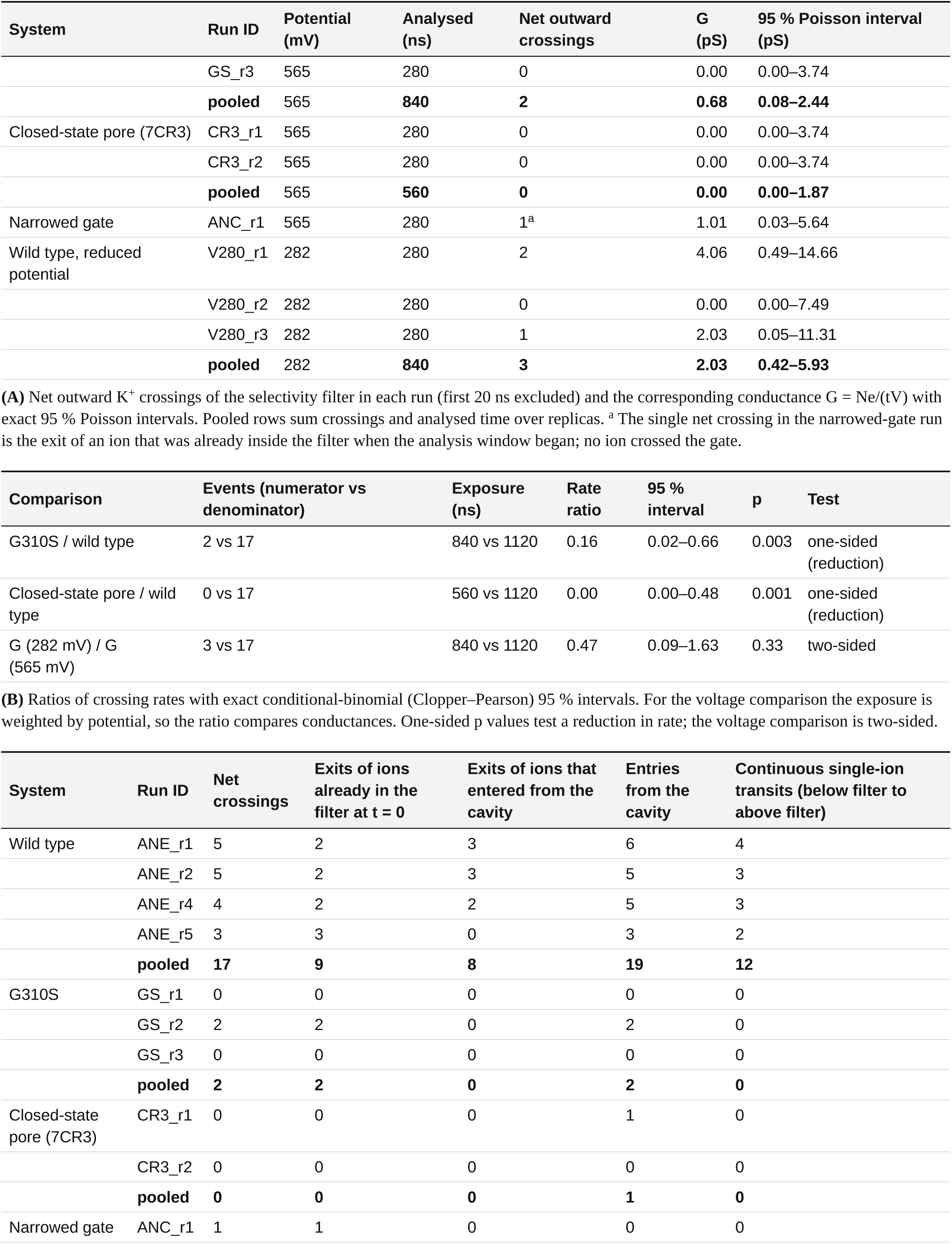
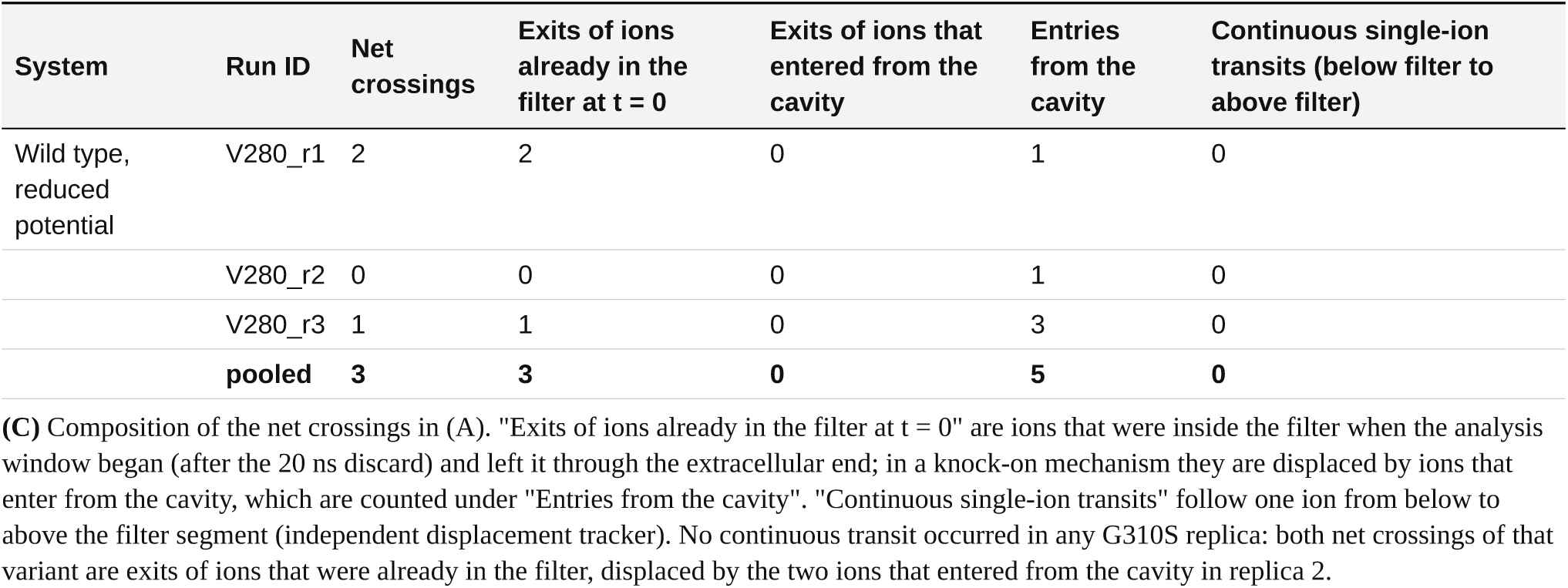
Net filter crossings and conductance.

**Table S3.** CHARMM36m control.

| System | Run ID | Window (ns) | Mean potential (mV) | Net transfer, filter exit | Net transfer, filter entrance (from cavity) | Cavity → filter entries | Pre-loaded exits | Displacement estimator (net ions) | Gate-flux estimator (net ions) |
| --- | --- | --- | --- | --- | --- | --- | --- | --- | --- |
| Wild type r1 | C36W_r1 | 280 | 574 | 1 | 0 | 0 | 1 | +0.06 | +0.00 |
| Wild type r2 | C36W_r2 | 280 | 576 | 1 | 0 | 0 | 1 | −0.30 | +1.00 |
| G310S r1 | C36G_r1 | 280 | 552 | 0 | 0 | 0 | 0 | +0.01 | +0.00 |
| G310S r2 | C36G_r2 | 280 | 561 | 0 | 0 | 1 | 0 | −0.02 | +0.00 |

| System | Run ID | Frames | d1 (Å) | Serine Oy-axis distance (Å) | Gate-plane water | Cavity water | K<sup>+</sup> in cavity (% frames) | K<sup>+</sup> below gate (% frames) |
| --- | --- | --- | --- | --- | --- | --- | --- | --- |
| Wild type r1 | C36W_r1 | 281 | 8.77 | – | 10.07 | 22.2 | 21.4 | 8.2 |
| Wild type r2 | C36W_r2 | 281 | 8.65 | – | 9.75 | 22.5 | 13.2 | 6.0 |
| G310S r1 | C36G_r1 | 281 | 8.49 | 3.44 | 4.52 | 22.8 | 0.0 | 6.8 |
| G310S r2 | C36G_r2 | 281 | 8.25 | 3.23 | 4.17 | 21.2 | 2.1 | 2.1 |

| Metric (mean ± SE of two replicas) | Wild type | G310S | Welch p (descriptive, n = 2 vs 2) |
| --- | --- | --- | --- |
| Gate-plane water | 9.91 ± 0.16 | 4.34 ± 0.17 | 0.002 |
| d1 (Å) | 8.71 ± 0.06 | 8.37 ± 0.12 | 0.17 |
| Cavity water | 22.35 ± 0.15 | 22.00 ± 0.80 | 0.74 |
| K<sup>+</sup> in cavity (% frames) | 17.30 ± 4.10 | 1.05 ± 1.05 | 0.14 |

| Metric (mean ± SE of two replicas) | Wild type | G310S | Welch p (descriptive, n = 2 vs 2) |
| --- | --- | --- | --- |
| K<sup>+</sup> below gate (% frames) | 7.10 ± 1.10 | 4.45 ± 2.35 | 0.45 |

| System | Net transfer at | Events / ns | G (pS) | 95 % interval (pS) | Expected at the Amber14sb rate | P(≤ observed), Poisson | p, conditional binomial (CHARMM36m < Amber14sb) |
| --- | --- | --- | --- | --- | --- | --- | --- |
| Wild type | filter exit | 2 / 560 | 1.00 | 0.12–3.60 | 8.5 | 9.3$\times 10^{-3}$ | 0.02 |
| Wild type | filter entrance (from cavity) | 0 / 560 | 0.00 | 0.00–1.84 | 8.5 | 2$\times 10^{-4}$ | 1$\times 10^{-3}$ |
| G310S | filter exit | 0 / 560 | 0.00 | 0.00–1.90 | 1.3 | 0.26 | 0.36 |
| G310S | filter entrance (from cavity) | 0 / 560 | 0.00 | 0.00–1.90 | 1.3 | 0.26 | 0.36 |

(D) Pooled CHARMM36m transfers compared with the number expected if CHARMM36m conducted at the Amber14sb rate over the same time (wild type 17 crossings in 1120 ns; G310S 2 in 840 ns): the Poisson probability of the observed count or fewer, and the conditional-binomial p for a rate below the Amber14sb rate.

**Table S4.** K^+^ coordination at the G310 plane.

| System | Run ID | Ion-frames in the gate plane | Water O | Serine Oy | Other protein O | Total | Bulk water O (control) |
| --- | --- | --- | --- | --- | --- | --- | --- |
| Wild type | ANE_r1 | 261 | 5.95 | 0.00 | 0.30 | 6.24 | 6.77 |
| Wild type | ANE_r2 | 350 | 6.03 | 0.00 | 0.22 | 6.25 | 6.78 |
| Wild type | ANE_r4 | 407 | 6.07 | 0.00 | 0.19 | 6.25 | 6.77 |
| Wild type | ANE_r5 | 248 | 6.31 | 0.00 | 0.13 | 6.44 | 6.78 |
| G310S | GS_r1 | 58 | 3.78 | 2.05 | 0.76 | 6.59 | 6.77 |
| G310S | GS_r2 | 221 | 4.00 | 1.73 | 0.65 | 6.38 | 6.78 |
| G310S | GS_r3 | 169 | 3.76 | 1.75 | 1.08 | 6.59 | 6.77 |

| First-shell oxygens per K<sup>+</sup> in the gate plane (replica means ± SE) | Wild type (n = 4) | G310S (n = 3) | Welch p |
| --- | --- | --- | --- |
| Water oxygens < 3.5 Å | 6.09 ± 0.08 | 3.85 ± 0.08 | < 10<sup>-4</sup> |
| Water oxygens < 3.0 Å | 5.06 ± 0.07 | 3.17 ± 0.03 | < 10<sup>-4</sup> |
| Serine Oy < 3.5 Å | 0.00 ± 0.00 | 1.84 ± 0.11 | 0.003 |
| Other protein oxygens < 3.5 Å | 0.21 ± 0.03 | 0.83 ± 0.13 | 0.03 |
| Total first-shell oxygens | 6.30 ± 0.05 | 6.52 ± 0.07 | 0.06 |
| Bulk water oxygens (control) | 6.77 ± 0.00 | 6.77 ± 0.00 | 0.72 |

(B) Replica means ± SE for wild type (four replicas) and G310S (three replicas) with Welch tests.

**Table S5.** Mixed tetramers (one and two variant subunits)

| System | Run ID | Analysed (ns) | Net outward crossings | of which exits of ions in the filter at t = 0 | of which exits of ions from the cavity | Continuous single-ion transits | Displacement estimator (net ions) | Gate-plane water (mean ± SD) | d1 (Å) | Serine Oy-axis (Å) | K<sup>+</sup> below the gate (% frames) |
| --- | --- | --- | --- | --- | --- | --- | --- | --- | --- | --- | --- |
| One serine subunit (H1) | H1_r1 | 280 | 2 | 2 | 0 | 0 | +2.00 | 7.89 ± 1.64 | 8.38 ± 0.39 | 4.91 | 17.4 |
| | H1_r2 | 280 | 5 | 2 | 3 | 3 | +4.97 | 7.78 ± 1.60 | 8.41 ± 0.36 | 4.46 | 14.9 |
|  | **pooled** | **560** | **7** |  |  |  |  |  |  |  |  |
| Two adjacent serine subunits (H2) | H2_r1 | 280 | 2 | 2 | 0 | 1 | +2.47 | 6.83 ± 1.69 | 8.24 ± 0.36 | 4.20 | 10.7 |
| | H2_r2 | 280 | 5 | 2 | 3 | 3 | +5.03 | 6.02 ± 1.54 | 8.19 ± 0.40 | 3.84 | 21.0 |
|  | **pooled** | **560** | **7** |  |  |  |  |  |  |  |  |

| System | Gate-plane water (replica mean) | f, replica SE (95 %) | f, 70 ns block means (95 %) | Additive expectation | Decision rule outcome (REJA §4) | Net crossings / ns |
| --- | --- | --- | --- | --- | --- | --- |
| H1 | 7.83 | 0.22 (0.13–0.32) | 0.22 (0.13–0.32) | 0.25 | additive | 7 / 560 |
| H2 | 6.43 | 0.58 (0.37–0.78) | 0.57 (0.47–0.67) | 0.50 | additive | 7 / 560 |

(B) The pre-specified read-out: $f = (W_{WT} - W_{mixed}) / (W_{WT} - W_{G310S})$ , computed from replica means ( $W_{WT} = 8.73 \pm 0.24$ , $n = 4$ ; $W_{G310S} = 4.73 \pm 0.12$ , $n = 3$ ) with delta-method intervals from replica standard errors and, in parallel, from 70 ns block means; the wider interval carries the decision. The additive expectation is $n/4$ for $n$ variant subunits. An independent recomputation using a different pore-axis definition and a different set of frames gave $f = 0.24$ (H1) and $0.60$ (H2).

**Table S6.** Excursions of ions towards the gate.

| System | Run ID | Analysed (ns) | Excursions from the vestibule | Reached the gate plane | Crossed to the cavity | Returned | Time in the plane per excursion, mean / median (ps) | K<sup>+</sup>-serine Oy contacts: number; median; longest (ps) |
| --- | --- | --- | --- | --- | --- | --- | --- | --- |
| Wild type | ANE_r1 | 280 | 369 | 30 | 6 | 24 | 86 / 40 | – |
|  | ANE_r2 | 280 | 287 | 22 | 6 | 16 | 154 / 105 | – |
|  | ANE_r4 | 280 | 382 | 27 | 5 | 22 | 148 / 50 | – |
|  | ANE_r5 | 280 | 303 | 9 | 3 | 6 | 274 / 120 | – |
| | **pooled** | **1120** | **1341** | **88** (79 per μs) | **20** (17.9 per μs; 23 %) | **68** | **141 / 60** | – |
| G310S | GS_r1 | 280 | 301 | 7 | 0 | 7 | 76 / 20 | 100; 35; 740 |
|  | GS_r2 | 280 | 216 | 17 | 2 | 15 | 130 / 30 | 240; 35; 1570 |
|  | GS_r3 | 280 | 231 | 14 | 0 | 14 | 121 / 15 | 133; 30; 1040 |
| | **pooled** | **840** | **748** | **38** (45 per μs) | **2** (2.4 per μs; 5 %) | **36** | **117 / 20** | **473; 30; 1570** |

| Depth reached, z* (nm from the G310 plane) | Wild type: excursions reaching z* (per μs) | of which crossed | fraction | G310S: excursions reaching z* (per μs) | of which crossed | fraction | Ratio of rates G310S / wild type |
| --- | --- | --- | --- | --- | --- | --- | --- |
| –0.50 | 758 (677) | 20 | 0.026 | 414 (493) | 2 | 0.005 | 0.73 |
| –0.40 | 355 (317) | 20 | 0.056 | 161 (192) | 2 | 0.012 | 0.60 |
| –0.30 | 143 (128) | 20 | 0.140 | 64 (76) | 2 | 0.031 | 0.60 |
| –0.25 | 89 (79) | 20 | 0.225 | 38 (45) | 2 | 0.053 | 0.57 |
| –0.20 | 61 (54) | 20 | 0.328 | 11 (13) | 2 | 0.182 | 0.24 |
| –0.10 | 43 (38) | 20 | 0.465 | 5 (6) | 2 | 0.400 | 0.16 |
| +0.00 | 32 (29) | 20 | 0.625 | 2 (2) | 2 | 1.000 | 0.08 |
| +0.10 | 21 (19) | 20 | 0.952 | 2 (2) | 2 | 1.000 | 0.13 |
| +0.25 | 20 (18) | 20 | 1.000 | 2 (2) | 2 | 1.000 | 0.13 |

(B) The same excursions resolved by the deepest position they reached: number of excursions whose maximum axial position was at least z\* (rate per microsecond in parentheses), how many of them went on to the cavity, and that fraction. The count at z\* = –0.25 nm differs by one from the gate-plane count in (A) because an ion can pass that level between two frames. Down to 0.25 nm below the plane the two systems differ by less than two-fold; the loss in G310S occurs between that level and the centre of the plane.

### Supplementary Figures

**Figure S1.**
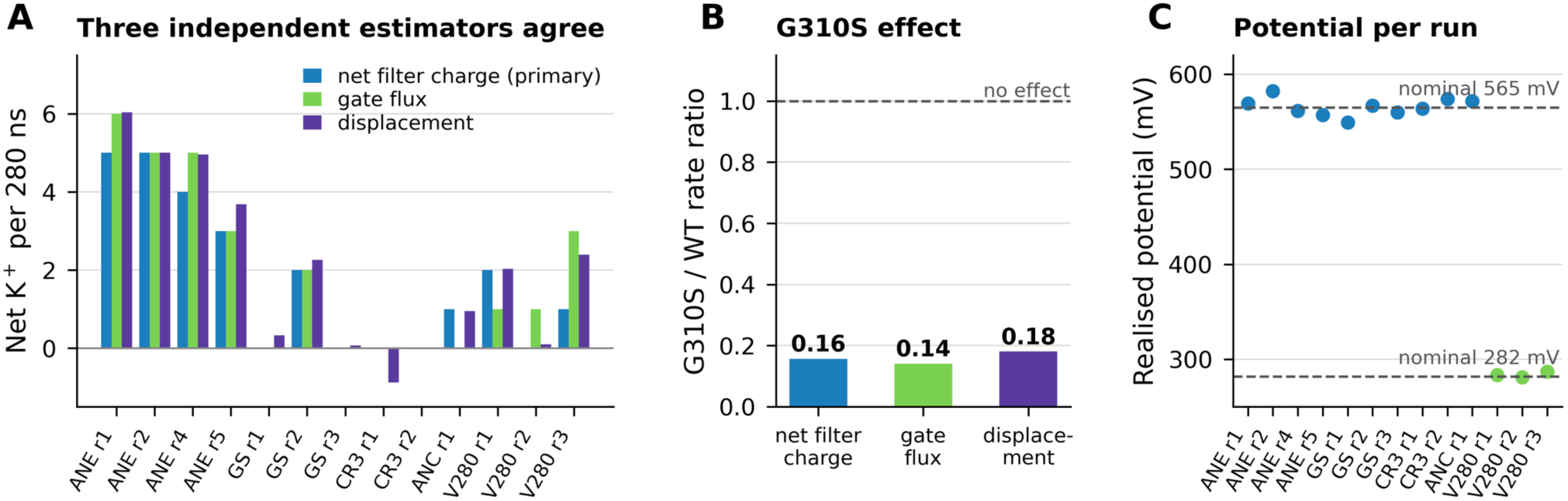
Three independent estimators agree. **(A)** Net K⁺ per 280 ns in every production run, counted three ways: net charge through the filter (the primary estimator, a zone-based state machine), flux through a slab at the gate, and a displacement integral over the filter segment. **(B)** The G310S/WT rate ratio is 0.16, 0.14 and 0.18 depending on the estimator. **(C)** The potential actually realised in each run, E·⟨L_z_⟩, against the nominal 565 and 282 mV.

**Figure S2.**
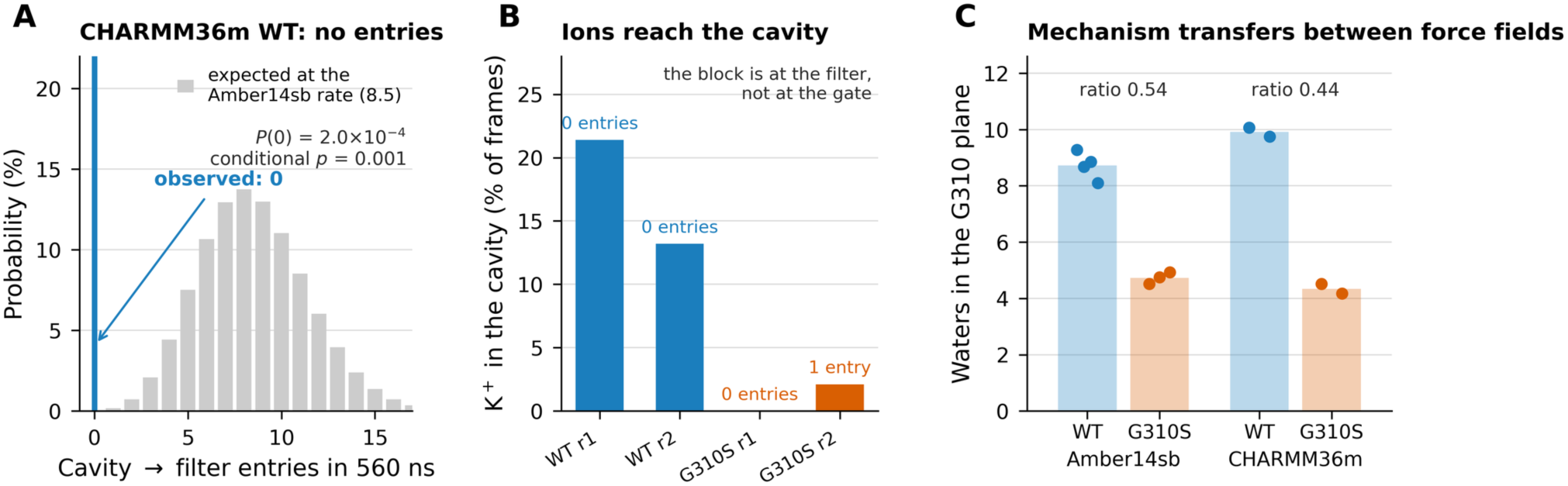
CHARMM36m control. **(A)** Distribution of cavity→filter entries expected in 560 ns at the Amber14sb rate (8.5 expected), against the zero observed with CHARMM36m. **(B)** The cavity is occupied by K⁺ in 13–21 % of wild-type frames, so ions do reach it; the block is at the filter. **(C)** The water column at G310 in both force fields: the wild-type/variant ratio is 0.54 in Amber14sb and 0.44 in CHARMM36m.

**Figure S3.**
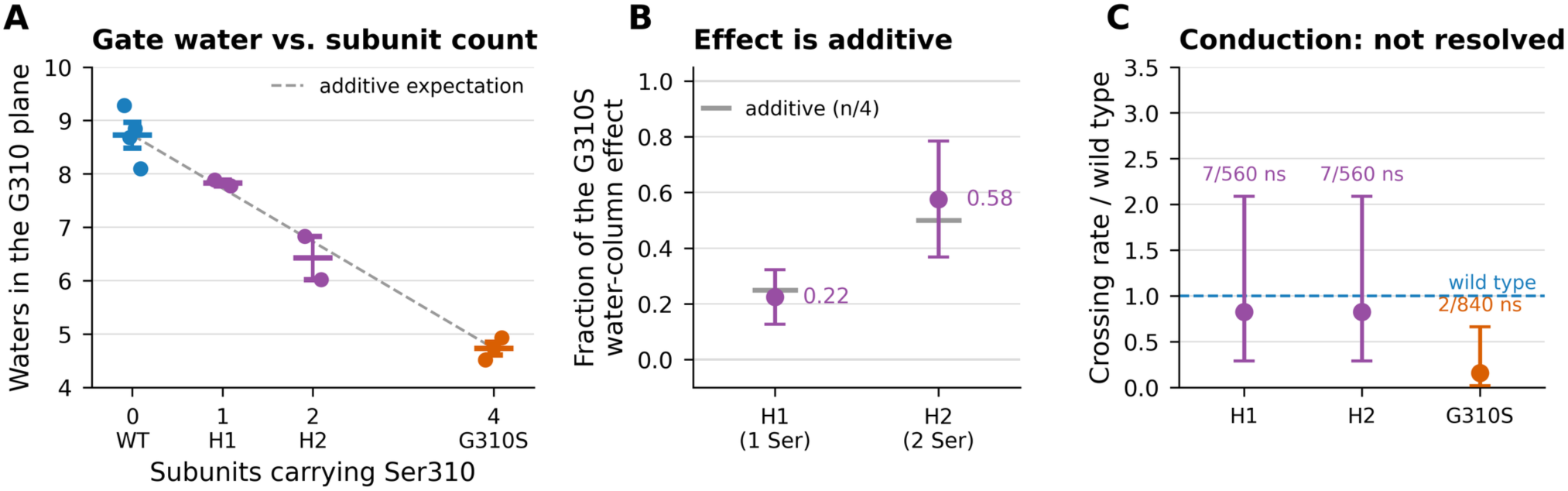
Mixed tetramers: the gate-water effect is additive. **(A)** Water molecules in the plane of G310 against the number of subunits carrying Ser310: wild type (four replicas), H1 with one variant subunit (two replicas), H2 with two adjacent variant subunits (two replicas) and G310S (three replicas). Points are replica means, the horizontal bars the group mean ± SE; the dashed line is the additive expectation drawn between the two homotetramers. **(B)** The same result as the fraction f of the G310S effect, with 95 % intervals from the delta method, against the additive expectation n/4 (grey bars). Both intervals contain the additive value. (C) Net crossing rate relative to the wild type, with exact 95 % conditional-binomial intervals; labels give net crossings and analysed time. With seven events per 560 ns the mixed pores are indistinguishable from the wild type and lie above G310S, so conduction does not resolve the question that the water column answers.

## Notes

https://github.com/Kushimboev/kcnq2-g310

### Competing Interest Statement

The authors have declared no competing interest.

